# A hybrid approach for automated mutation annotation of the extended human mutation landscape in scientific literature

**DOI:** 10.1101/363473

**Authors:** Antonio Jimeno Yepes, Andrew MacKinlay, Natalie Gunn, Christine Schieber, Noel Faux, Matthew Downton, Benjamin Goudey, Richard L. Martin

## Abstract

As the cost of DNA sequencing continues to fall, an increasing amount of information on human genetic variation is being produced that could help progress precision medicine. However, information about such mutations is typically first made available in the scientific literature, and is then later manually curated into more standardized genomic databases. This curation process is expensive, time-consuming and many variants do not end up being fully curated, if at all. Detecting mutations in the literature is the first key step towards automating this process. However, most of the current methods have focused on identifying mutations that follow existing nomenclatures. In this work, we show that there is a large number of mutations that are missed by using this standard approach. Furthermore, we implement the first mutation annotator to cover an extended mutation landscape, and we show that its F1 performance is the same performance as human annotation (F1 78.29 for manual annotation vs F1 79.56 for automatic annotation).

## Introduction

As the cost of DNA sequencing continues to fall, an increasing amount of information on human genetic variation is being produced that could help progress precision medicine. Increasingly, improved understanding of an individual’s genetic mutations is being used to identify individuals at high risk for a given disease, make better predictions about disease prognosis and tailor treatments so that they best suit the given patient. New findings about genetic mutations typically first appear in the scientific literature. This novel information is then later manually curated into genomic databases, such as COSMIC (Catalogue of Somatic Mutations in Cancer)^1^ where the information is presented in a standardized, queryable format. While such databases are invaluable resources, this curation process is expensive, time-consuming and only captures a small part of the relevant literature^2^. Furthermore, in addition to the scientific literature, large collections of health records with mutation information can be considered as a complementary source.

Identification of mutations in the scientific literature (often referred to as ‘mutation mentions’) has been covered in previous work, including comparisons of existing methods^2^. Most existing methods for mutation annotation have focused only on identifying mutation mentions that are related to changes in DNA/protein sequences and are similar to standardize nomenclature^3-5^. However, this may miss other types of mutations, such as those in RNA or with database-specific identifiers, as well as mentions of mutations without specific DNA/protein change (e.g. mutation, translocation, deletion, etc. close to specific gene identifiers). Such approaches also ignore a large set of complementary entities, e.g. epigenetic alterations, micro-RNAs, that are critical for improving personalized medicine. For instance, the following snippet from PubMed citation PMID:15993480 highlights epigenetic related methylation that would be otherwise missed: “**promoter hypermethylation** is not the major cause of decreased expression of **TbetaRII** in **endometrial cancers**”. This variety of entities implies as well that a single methodology for the identification of these entities in the scientific literature might not be sufficient.

In this paper, we take a broad view of the human mutational landscape contained in the scientific literature and explore a method for automatically identifying such mutations. We provide empirical evidence showing that a large number of mutations are missed by existing methods that consider only DNA mutation similar to those in standardized nomenclature. We overcome these limitations by extending what has been considered for mutation annotation to cover a wider range of possible mentions of genetic variations in the scientific literature. This corpus is available from (https://github.com/ibm-aur-nlp/amia-18-mutation-corpus). A hybrid approach to mutation annotation is proposed, in which both rule-based and machine learning methods are combined to cover the broad mutation landscape. Two machine learning methods are compared: a deep learning approach of a bidirectional LSTM (Long Short Term Memory) with a neural CRF (Conditional Random Field) compared to a standard CRF approach with an engineered feature set. The performance of the hybrid developed approach incorporating our deep learning method achieves the same F1 performance as human annotation (F1 78.29 for manual annotation vs F1 79.56 for automatic annotation). This is the first implementation of a mutation tagger that covers an extended mutation landscape and provides a strong foundation for tagging other entities critical for improving precision medicine.

## Methods

In this section, we describe the annotation guidelines and the manual annotation process, which includes an analysis of the inter-annotator agreement. Then, we describe the automatic annotation method for mutation annotation. This method has been implemented using UIMA (Unstructured Information Management Architecture)^6^ (https://uima.apache.org). On top of UIMA we have used uimaFit (https://uima.apache.org/uimafit.html) and ClearTK^7^ (https://cleartk.github.io/cleartk) for machine learning algorithms.

### Annotation guidelines

The domain of interest in our work is colorectal cancer, which was used to help define a subset of citations from MEDLINE^1®^ for annotation. MEDLINE is the largest biomedical bibliographic database and it currently contains more than 28 million citations. Initially, we manually selected a set of 10 citations from PubMed^®^ that were examples of the domain of interest, and used these to generate a PubMed query in order to identify relevant articles. The query that we generated is “((colorectal cancer) AND Humans[MH]) AND (genetic variation[MH] OR mutations)”, which recovered around 20k citations from PubMed.

Prior to the discussion with the annotators, we briefly defined what should be included as entities for annotation, which initially consisted of mutations and gene/protein mentions. Similar to previous work^5^, we found that it is difficult to make the distinction between genes and proteins in a MEDLINE citation, so the entity *Gene/protein* was defined. Before the annotation process, we used two batches of 10 MEDLINE citations to calibrate the guidelines. During this calibration process, annotators were asked to follow the initial guidelines while annotating 10 citations and disagreements were discussed afterwards. Due to the complexity of the biomedical domain, and to have a complete annotation of possible sequence modification mentions, additional entities, such as genomic locations and other nonmutation modifications (e.g. epigenetic alterations), were identified. These additional entities helped the annotators to categorize all possible modification mentions in MEDLINE citations. All these modifications considered for manual annotation have possible phenotypic implications that have not been considered in this work. After the calibration process, the following entities were defined:

**Mutation:** Any term denoting a mutation, which includes HGVS nomenclature mutations. Mutations are further divided into *DNA mutation, Protein mutation, RNA mutation* and *dbSNP* database^8^ (https://www.ncbi.nlm.nih.gov/SNP) identifiers. These sub-types have been merged into the mutation entity.

**DNA modification:** In addition to mutations, there are other non-sequence-altering changes that may produce over or under expression of a gene product. Mentions of these changes in text were annotated as DNA modifications. Examples include epigenetic alterations. Identifying DNA modifications supports the understanding of factors that affect gene expression.

**Gene/protein:** Mentions of specific genes/proteins are annotated (e.g *BRCA1*). Generic words such as *gene* or *protein* have not been annotated. NCBI’s Gene^9^ (https://www.ncbi.nlm.nih.gov/gene) and Uniprot^10^ (http://www.uniprot.org) were considered as reference resources during annotation.

**Locus:** Position on a chromosome or protein, e.g. *nucleotide 986-1218* or CPG island.

**RNA:** Mutations can be identified as well in RNAs (ribonucleic acid) and were annotated as an entity. This entity type includes mentions of micro-RNAs.

In addition to entities, several relations between the entities were defined, which had to appear in the same sentence:

**Component of:** Relate a locus that is a *component of* a gene.

**Has modification:** Relation between gene/protein and DNA modification.

**Has mutation:** Relation between Gene/protein/RNA/locus entities and mutation entities.

### Manual corpus annotation

Annotation was performed by 5 domain experts using the BRAT tool^11^ (http://brat.nlplab.org) (e.g. figure 1) on the abstract and title of selected MEDLINE citations. MEDLINE citations were first separately annotated by two experts. Disagreements between the pair of annotators were then resolved by the pair working on the same citation jointly. We have found that by doing this annotation procedure, the quality of the manual annotation improved significantly and while there are cases in which the annotators disagree due to the complexity of the domain, we realized that recall was largely improved.

**Figure 1:**
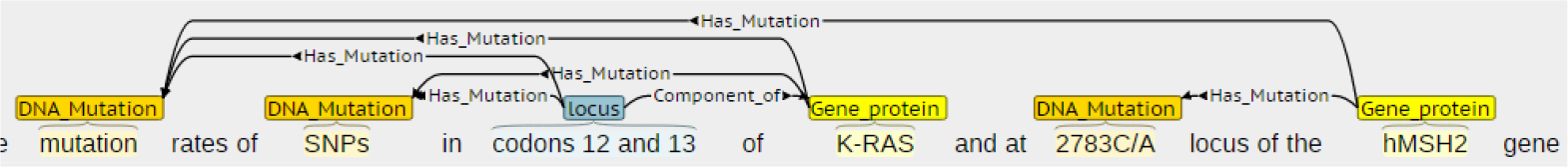
Example BRAT annotation

To determine inter-annotator agreement rates, a set of citations was annotated by four annotators similar to the method defined above. Each document was first annotated by subject matter experts, disagreements were then resolved between a pair of annotators. Inter-annotator agreement was measured using the documents resolved by one pair versus the documents resolved by the other pair. Table 1 shows the inter-annotator agreement in our manually annotated set measured using precision, recall and F1^5^, using the standard formulas (TP = True Positives, FP = False Positives, FN = False Negatives, *Precision* = *TP*/(*TP* + *FP*), *Recall* = *TP*/(*TP* + *FN*), *F*1 = (2 * *Precision* * *Recall*)/(*Precision* + *Recall*)). Precision, recall and F1 are also used in the Results section to evaluate the automatic mutation annotators.

**Table 1:** Inter-Annotator Agreement from our manually annotated set

| Entity | Exact |  |  | Partial |  |  |
| --- | --- | --- | --- | --- | --- | --- |
|  | Precision | Recall | F1 | Precision | Recall | F1 |
| **DNA modification** | 64.71 | 78.57 | 70.97 | 76.47 | 92.86 | 83.87 |
| **Gene/protein** | 93.42 | 88.36 | 90.82 | 96.35 | 91.55 | 93.89 |
| **Locus** | 62.67 | 83.42 | 71.57 | 70.22 | 92.94 | 80.00 |
| **Mutation** | 79.03 | 77.57 | 78.29 | 87.97 | 87.39 | 87.68 |
| **Component of** | 41.86 | 78.26 | 54.55 | 48.83 | 80.77 | 60.87 |
| **Has modification** | 44.00 | 100.00 | 61.11 | 56.00 | 100.00 | 71.79 |
| **Has mutation** | 54.14 | 69.23 | 59.86 | 70.69 | 72.44 | 71.55 |

Inter-annotator agreement was evaluated using two different methods to define matching annotations: one that compares the exact boundaries of the span of text (Exact matching evaluation) and another one that relaxes this to any overlapping span (Partial matching evaluation). Partial matching is interesting since it provides insights about the complexity of identifying the boundaries of the entities.

Comparison of corpora for evaluation was performed using the brateval tool (https://bitbucket.org/nicta_biomed/brateval). For relation annotation agreement, we take into account the identification of the entities that take part in the relation and the relation between them.

There is high agreement in gene/protein and mutation annotation, as in previous work^5^. When boundaries are relaxed the agreement improves, which shows that annotators agree in the location of the entities’ spans.

The final data set contains 167 MEDLINE citations, which we split into 122 citations for training our automatic mutation annotators, and a further 45 for testing their performance. According to the identified annotations, the corpus contains 60 mentions of *DNA modifications*, 1324 mentions of *genes/proteins*, 320 mentions of *loci*, 1337 mentions of *mutations* and 23 mentions of *RNAs*. The expert annotators also identified 94 *component of* relations, 52 *has modification* relations and 907 *has mutation* relations. Table 2 shows the most frequent terms by type.

**Table 2:**
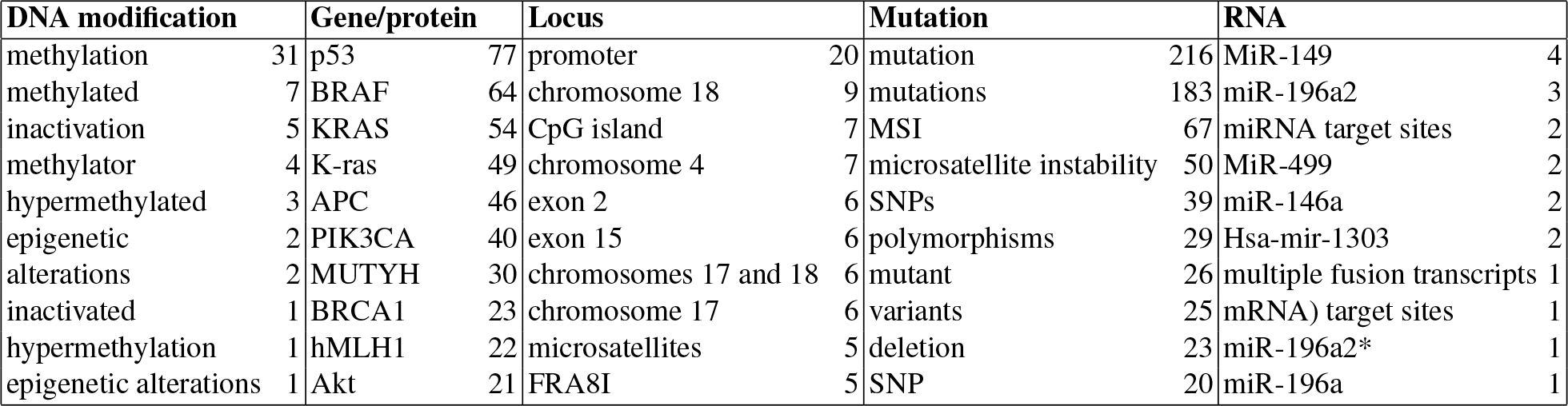
Top 10 most frequent terms per entity type in our manually annotated corpus

### Regular expression annotation

For human mutations, a standardized nomenclature has been defined by the Human Genome Variation Society (HGVS) (http://www.hgvs.org/mutnomen). This nomenclature covers RNA, cDNA and protein mutations. Mutations can also follow nomenclature defined by the International System for Human Cytogenetic Nomenclature (ISCN)^12^ (https://www.hgvs.org/mutnomen/ISCN.html) or the Human Cytochrome P450 (CYP) Allele Nomenclature (https://www.pharmvar.org/htdocs/archive/index_original.htm). Some databases, such as dbSNP, also have their own unique identifiers.

We have developed a set of regular expressions that can match these standard sets of nomenclature in text. These expressions allow identifying these mutations in text and, since we can identify the expression that has triggered the annotation, we can identify the type of mutation, e.g. substitution, deletion, insertion. Similar systems that rely on regular expressions have been previously developed including methods for detecting protein point mutations^4^ and broad ranges of DNA mutations^13^. Separate sets of regular expressions were developed to identify nomenclature by HGVS, ISCN and CYP. Regular expressions have also been prepared to identify dbSNP ids (*rs*[0 – 9]+).

Unfortunately, authors do not always exactly follow the HGVS or ISCN nomenclature. The most common differences in the HGVS nomenclature consists in not using the prefix for gene (*c*.) and protein (*p*.) mutations. Additional examples include adding spaces in the mutation term or not using the characters specified. For example, the cDNA mutation *2783C/A* in MEDLINE citation PMID:25561800 should have been written as *c.2783C>A*. We have developed additional expressions to identify the most common cases of nomenclature abuse and provided a method that creates a canonical HGVS mutation form. The training set has been used to develop and tune the regular expressions.

### Dictionary-based annotation

We derived a dictionary of terms from our training data, excluding mutation entities identified by our regular expressions, that directly identity specific types of entities. Extracted terms, which are typically multi-word expressions (e.g. *single nucleotide polymorphism*, *genomic aberration*, *microsatellite heteroduplex*), denote a mutation in our training set and it is assumed that no context is required for their annotation. This dictionary was used to annotate mutations in text wherever an entry was observed. As shown in the results, this dictionary approach has a limited recall but complements the annotations of the other methods.

### CRF-based annotation

CRF^14^ has been widely used for named entity recognition, including the biomedical domain^15,16^. In our work, we have used a linear chain CRF implementation provided by CRFSuite^17^. We have used the following engineered feature set: (a) surface form of the token, (b) information about upper/lower case character in the token, (c) if the token is a number or not, (d) tokens neighbouring the current token being annotated and (e) a flag to indicate if the token has been found in a mutation dictionary we developed, based on extracting terms from the MeSH^®^ controlled vocabulary (https://meshb.nlm.nih.gov/search) tree branches *Mutation (id:D009154*) and *SNP (id:D011110*).

### Deep learning-based annotation

In addition to the CRF annotation-based method, we have evaluated a deep learning method that we have developed for named entity recognition^18^. This method, as shown in figure 2, is comprised of an encoding mechanism that uses a bidirectional LSTM network (biRNN)^19^, which acts as a feature encoding that substitutes manual engineering of features, and a decoding method that uses a neural CRF^20^ to generate the named entity recognition annotation tags. Three layers of biRNN have been stacked using residual connections. In the biRNN the LSTM dimension is 25 since our data set is smaller compared to previous work, which was 100. All the other parameters are the same as in previous work^18^.

**Figure 2:**
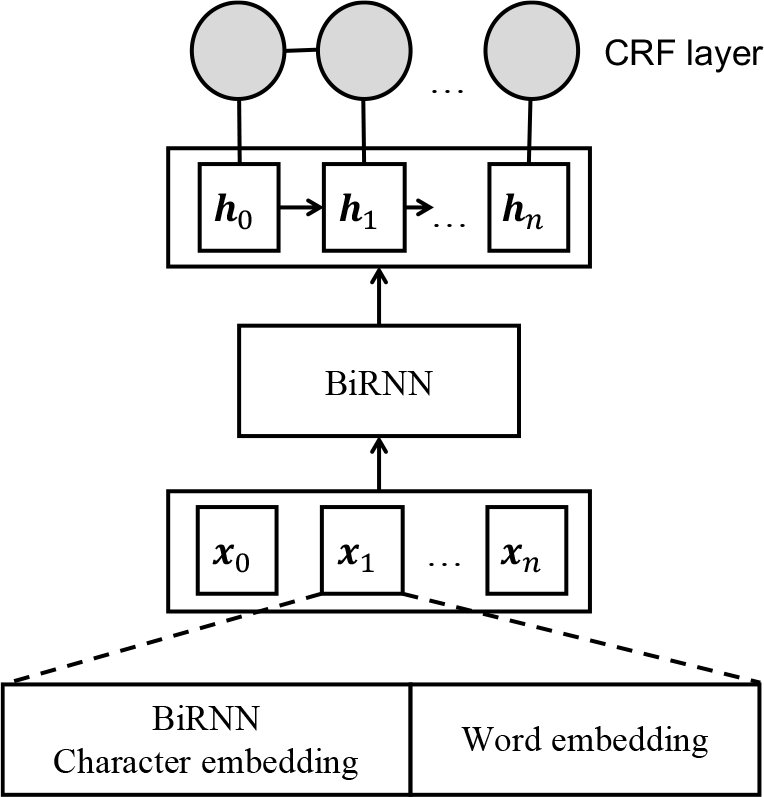
Named entity recognition system based on a bidirectional LSTM encoder with neural CRF as decoder

Input features are word embeddings, and character embeddings derived from the training set tokens and no pretraining was done for character embeddings. The corpus for the generation of the word embedding is comprised of MEDLINE 2017 and PubMed Central^®^ Open Access journal articles obtained from (https://www.ncbi.nlm.nih.gov/pmc/tools/openftlist). Embeddings were generated using the word2vec tool^21^ (https://code.google.com/archive/p/word2vec). Dimension of the word embedding vectors is 100 with a window of 5 words. Skip-gram was used as the method for preparing the word embeddings. No additional features have been engineered.

### Hybrid combination of mutation annotation methods

We have used several methods to cover different types of mutations. A regular expression-based system is particularly well-suited for extraction of a certain kind of phrase, such as a HGVS nomenclature, which conforms to a well-defined pattern. A dictionary-based method performs well for extracting phrases that are very likely to be a mutation no matter what the context is that they occur in, such as *single nucleotide polymorphism*. In contrast to these two approaches, a machine learning-based system is particularly suited for mentions whose variety is too great to be captured exhaustively in a dictionary, and which can also not be defined by a precise systematic phrasing; here it might be a better choice to use a model which can *learn* to detect these phrases based on the context of the mention.

By performing annotation using a combination of these approaches, taking the best properties of each method, we aim to better approximate human parsing of text. A human reader would be able to identify some phrases in text as being mutations because they are established phrases (analogous to a dictionary-based method), some because they follow a pattern that the expert is aware of (analogous to a regular expression-based method), and others still by interpreting the mention and its structure within the particular context it occurred in (machine learning-based method).

An outline of our hybrid approach is shown in figure 3. To merge the annotations, the intention is to consider all annotations by each individual methods and merge them by selecting the maximum span. During conflict resolution, all nested annotations are removed.

**Figure 3:**
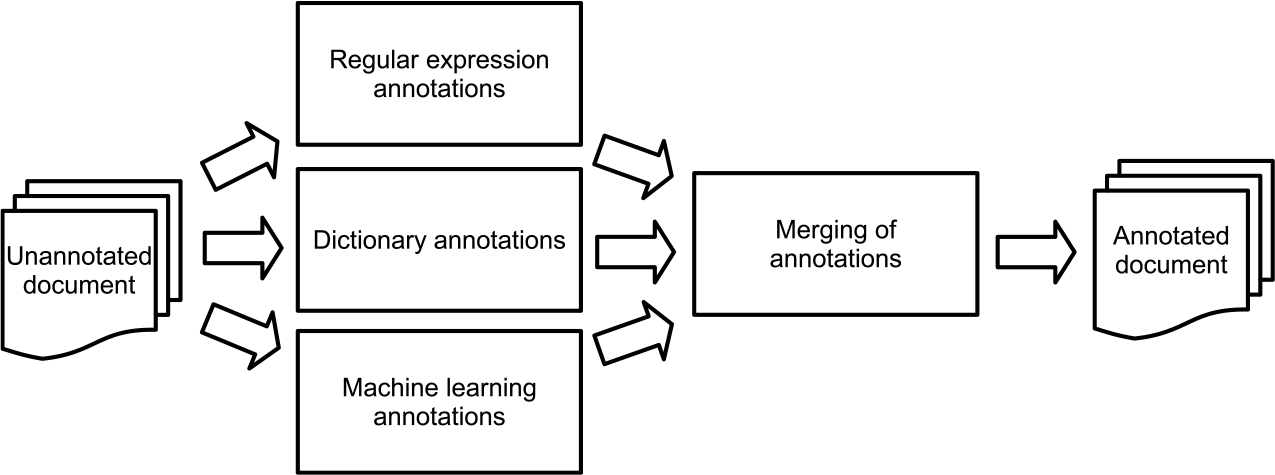
Mutation annotation pipeline. Machine learning annotations are provided by the CRF or the neural method.

## Results

Results for the four individual methods and the hybrid method presented above are shown in Table 3. Performance is shown in terms of precision, recall and F1 measure (defined in the Methods section) across both exact and partial matching evaluation. The two types of evaluation show some differences across all methods with partial matching yielding slight-moderate increases in performance across all three metrics for most methods. This is expected given its less stringent criteria.

**Table 3:** Automatic mutation annotation results

| Method | Exact |  |  | Partial |  |  |
| --- | --- | --- | --- | --- | --- | --- |
|  | Precision | Recall | F1 | Precision | Recall | F1 |
| Inter Annotator Agreement | 79.03 | 77.57 | 78.29 | 87.97 | 87.39 | 87.68 |
| Regular Expressions (RegEx) | **93.75** | 3.90 | 7.48 | 93.75 | 3.90 | 7.48 |
| Dictionary Annotation (DA) | 81.51 | 25.19 | 38.49 | 92.44 | 28.57 | 43.65 |
| Conditional Random Field (CRF) | 83.71 | 57.40 | 68.10 | 93.56 | 64.32 | 76.23 |
| Deep learning (DL) | 82.04 | 68.83 | 74.86 | 91.64 | 77.08 | 83.73 |
| RegEx + CRF + DA | 82.02 | 67.53 | 74.07 | 91.80 | 75.78 | 83.02 |
| RegEx + DL + DA | 84.96 | **74.81** | **79.56** | **93.81** | **82.81** | **87.97** |

The different automatic mutation annotators also showed considerable differences. Regular expressions (Regex) show high precision, comparable to previous work, but with lower recall compared to the total number of mutation mentions available. This is expected since they only cover mentions of mutations in HGVS format and the supported variants, which are only some of the mutations in the data set (less than 4%, the Regex method has a recall of over 93% when considering only HGVS and nomenclature based mentions in our manual set, which is similar to existing methods^2^).

Two machine learning methods were evaluated: a CRF with engineered features and a deep-learning approach. Under both exact and partial matching, the deep learning method performs better compared to the CRF approach. Since the deep learning method relies on a neural CRF, the improvement in performance might be due to the improved representation obtained by the encoding biRNN supported by the pretrained word embeddings. The hybrid approach, either based on CRF or deep learning, improves the performance of individual methods. In particular, the hybrid approach obtains similar F1 performance compared to the inter-annotator agreement. Looking at the recall of the combined methods, the dictionary method (DA) seems to complement the regular expressions and machine learning methods.

## Discussion

Identifying a broader set of entities that should be considered. In our manual annotation, we have considered mentions of mutations without specific DNA/protein change (e.g. mutation, translocation, deletion, …). These terms provide additional information about the potential changes happening in a gene that were not available in previous annotations. The additional mutations complement the information about mutations expressed in standardised HGVS format or similar, which is only a small portion of our manual annotation.

To predict this broader range of mutations, we proposed a hybrid framework using a combination of rule-based and machine learning-based approaches. The performance of the hybrid mutation annotator, making use of our previously proposed deep learning method, achieves similar performance to the inter-annotator agreement in terms of F1. Our regular expressions have high precision but achieve a recall of 3.90%. This is expected since there is small portion of mutations consider in our data set that follow a specific nomenclature, e.g. HGVS.

To better understand how these methods complement each other we show some examples. Citation PMID:17990317 contains mentions of HGVS nomenclature mutations missed by the other methods. Certain variations of terms such as *mutant* or rarer terms such as *aneuploidy* are complemented by dictionary terms. Machine learning methods use the context to better contextualize words. The deep learning method is able of identifying better multi-word expressions (e.g. genetic variation vs variation by the CRF method) and is able of identifying additional expressions (e.g. *AAC to AAT* in PMID:10690536).

There are terms missed by the hybrid annotator such as *allelic differences* or variations of the term mutation, such as *mutational*, or microsatellite instability such as *microsatellite unstable*. There are some boundaries differences with terms such as *DNA aneuploidy* vs *aneuploidy*, which seem to be difficult to characterize as well in the manual annotation. Another example would be *monoallelic and biallelic mutations* vs *biallelic mutations*.

When using the mutation annotator across all MEDLINE citations, we identify a total of 6,459,550 mentions of mutations. The 10 most frequent terms already make 3,883,830 mentions of *mutations*, which would have been missed by existing methods. As most of the mutations mentions link back to a specific gene, this is strong evidence that the proposed mutation annotator is likely to capture a significant amount of information ignored by existing methods.

## Conclusions and Future Work

In this work, we move beyond the tagging of only DNA/protein mutations that appear similar to standardized nomenclature, to try and tag a much larger range of variation available in the scientific literature. We have provided experimental results in the annotation of mutations from the scientific literature and shown that a hybrid method using deep learning obtains improved predictive performance beyond the individual baseline methods.

In future work, we would like to extend the current named entity recognition approach to the other entity types such as locus, or DNA modifications considered in the manual annotation. This would include automatic annotations of relations as annotated in the manual annotation. We have not considered the annotation of the phenotypic consequences of mutations that will be included in future work, extending the current manual annotations in our corpus.

